# Mouse CHD4-NURD is required for neonatal spermatogonia survival and normal gonad development

**DOI:** 10.1101/687137

**Authors:** Rodrigo O. de Castro, Luciana Previato, Agustin Carbajal, Victor Goitea, Courtney T. Griffin, Roberto J. Pezza

**Author notes:** Corresponding author: Roberto J. Pezza. Suite B305. 825 NE 13^th^ street, Oklahoma City, Oklahoma, 73104.

## Abstract

Testis development and sustained germ cell production in adults rely on the establishment and maintenance of spermatogonia stem cells and their proper differentiation into spermatocytes. Chromatin remodeling complexes regulate critical processes during gamete development by restricting or promoting accessibility of DNA repair and gene expression machineries to the chromatin. Here, we investigated the role of CHD4 and CHD3 catalytic subunits of the NURD complex during spermatogenesis. Germ cell-specific deletion of Chd4 early in gametogenesis, but not Chd3, resulted in arrested early gamete development due to failed cell survival of neonate undifferentiated spermatogonia stem cell population. Candidate assessment revealed that CHD4 controls expression of Dmrt1 and its downstream target Plzf, both described as prominent regulators of spermatogonia stem cell maintenance. Our results show the requirement of CHD4 in mammalian gametogenesis pointing to functions in gene expression early in the process.

## Introduction

Defects in gametogenesis are a leading cause of infertility and an important cause of birth defects associated with aneuploidy. Insights into the mechanisms underlying testis formation, including spermatogonia stem cell, are necessary to improve the outcomes of common gonad developmental diseases.

In mice, spermatogenesis begins from isolated germ cells called spermatogonia A-singles (As) that undergo to a series of mitotic divisions to produce spermatogonia known as paired (Apr) and aligned (Aal) that contains chains of 4 to 16 cells anchored by intercellular bridges as a result of incomplete cytokinesis [1, 2]. At this point, spermatogonia cells start a process of differentiation (A1, A2, A3, A4 or intermediate (In)), formation of type B spermatogonia, and then transition to pre-leptotene cells that initiate the series of meiotic divisions that ultimately originate spermatozoa.

Chromatin undergoes extensive remodeling during gametogenesis, leading to altered gene expression and chromosome organization, and ultimately controlling obligatory developmental transitions such as the conversion from undifferentiated to differentiated spermatogonia, spermatogonia commitment to meiosis, and meiotic progression [3, 4]. The NURD (NUcleosome Remodeling and Deacetylase) is a prominent chromatin modifying complex that functions to control gene expression via chromatin remodeling and histone deacetylation [5, 6]. The NURD complex contains two highly conserved and widely expressed catalytic subunits, CHD3/Mi-2α (chromodomain-helicase-DNA-binding 3) and CHD4/Mi-2β, which are members of the SNF2 superfamily of ATPases [5–9]. NURD plays a central role in various developmental and cellular events, such as controlling the differentiation of stem cells, maintaining cell identity, and responding to DNA damage [9–11]. In testis, CHD5 is required for normal spermiogenesis and proper spermatid chromatin condensation [12], while CHD3/4 has been described to localize at the X-Y pseudoautosomal region, the X centromeric region, and then spreads into the XY body chromatin [13, 14]. Although the role of CHD5 has been well defined, no studies to date have addressed the requirements or mechanisms of CHD4 and CHD3 in any germ cell type.

In this study, we report that CHD4 (but not CHD3) is essential for testis development and sustained germ cell production. Germ cell-specific deletion of CHD4 results in the developmental arrest of undifferentiated spermatogonia in neonatal mice progressing to a Sertoli-only phenotype. Our studies of selected CHD4 target genes and subsequent cytological and expression analysis show that CHD4 control Dmrt1 gene expression and downstream targets such as Plzf. We propose these results suggest a possible mechanism by which CHD4 contributes to early germ cell development regulating genes that are required for survival/maintenance of spermatogonia cells.

## Results

### *Chd4* expression during mouse germ cell development

To investigate a potential role for CHD4 in germ cells, we assessed CHD4 expression in newborn and adult mouse testes by immunofluorescence. CHD4 was highly expressed in spermatogonia cells (marked by PLZF, aka ZBTB16) and in Sertoli cells (marked by SOX9) (Fig. 1A and Fig. S1) but not in the negative control (PLZF-positive spermatogonia from *Ddx4-Chd4^-/-^* testes, Fig. S2A). PLZF is expressed in undifferentiated spermatogonia [15, 16] (Fig 1A). In agreement with a previous report [13], immunosignal of CHD4 was detected in late-pachytene stages (Fig. 1A, selected area on adult mouse top panel). In sum, CHD4 is detected in spermatogonia, Sertoli cells, and primary spermatocytes.

**Figure 1.**
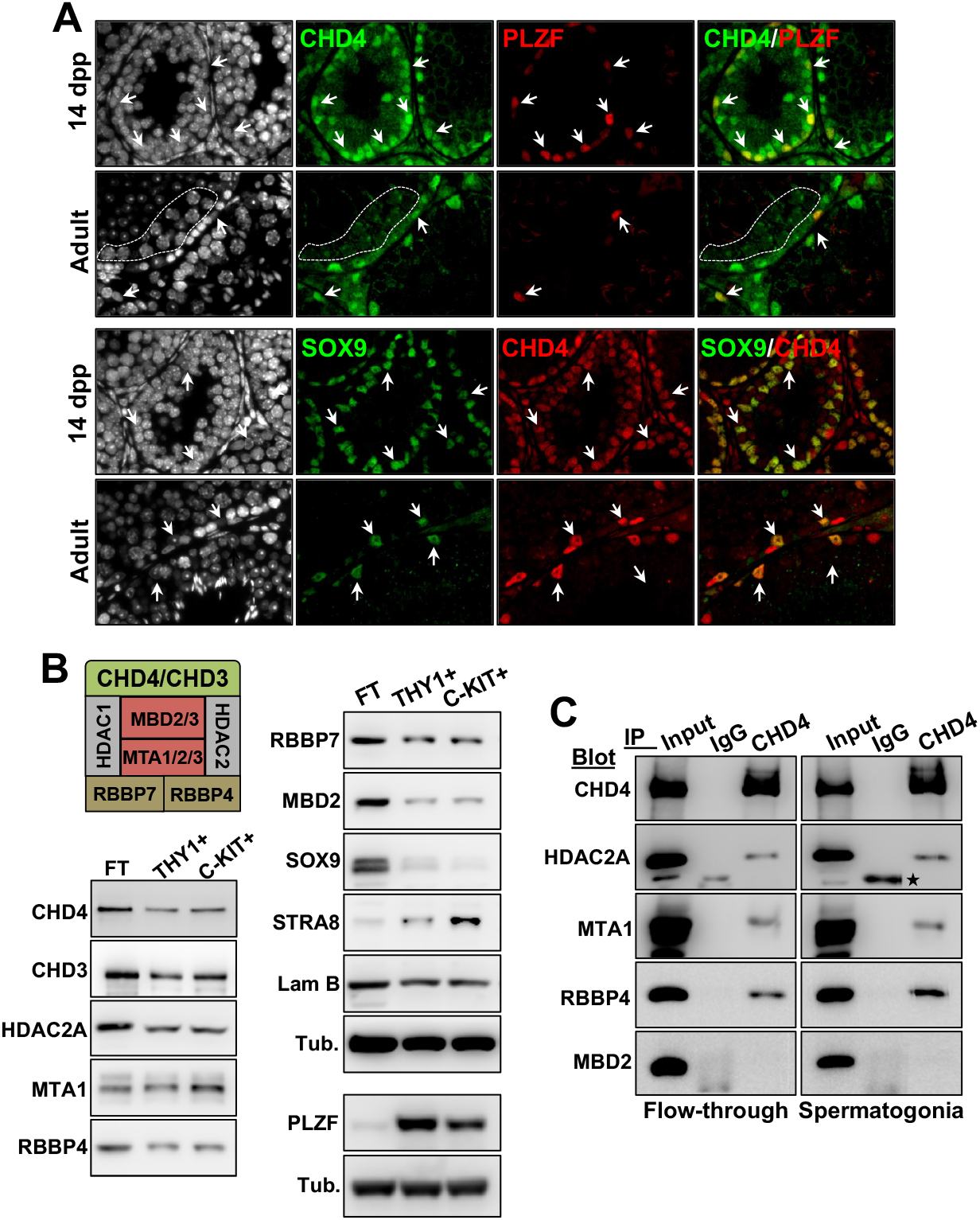
CHD4 expression and formation of different NURD complexes during gametogenesis. **(A)** Expression of *Chd4* monitored by immunofluorescence in testis sections of 14 dpp and 2 months (adult) old mouse using two distinct antibodies against CHD4, a rabbit polyclonal anti-CHD4 and anti-PLZF (top panel) and mouse monoclonal anti-CHD4 and anti-SOX9 (bottom panel). Arrows indicate examples of CHD4 positive spermatogonia (PLZF) or Sertoli (SOX9) cells. Cells within the punctuated line are pachytene spermatocytes. Similar results were observed using three different mice. **(B)** Western blot analysis of different NURD components in samples of cells enriched in undifferentiated spermatogonia (THY1.2+) (PLZF positive), differentiating spermatogonia (c-KIT+) (STRA8 positive) and the flow-through (FT), the latter from the spermatogonia enrichment procedure, which contains mostly somatic cells, including a large amount of Sertoli cells (SOX9 positive). Lamin B (Lam B) and α-tubulin (Tub.) were used as loading standards. The scheme represents the composition of a canonical NURD complex. **(C)** CHD4-participating NURD complexes in enriched fractions of spermatogonia or cells in the flow-through assessed by co-immunoprecipitation with anti-CHD4 as a bait. Nonspecific IgG was used as a bait control. Experiments were repeated twice, and the star marks an unspecific reactive band.

We confirmed that CHD4 is expressed at pre-meiotic stages of male gamete development by analyzing CHD4 protein levels by western blot in enriched fractions (see methods for details) of undifferentiated and differentiating spermatogonia (obtained from wild type 7dpp mice and using THY1.2+ and c-KIT+ affinity columns, respectively) (Fig. 1B). CHD4 was also detected in the affinity column flowthrough (which was enriched mostly in Sertoli cells, here demonstrated by immunoblotting of Sox9). The level of cell population enrichment was assessed by western blot and markers specific for undifferentiated spermatogonia (PLZF) and differentiating spermatogonia (STRA8) (Fig. 1B). Additionally, we analyzed cell fractions enrichment by immunofluorescence (Fig. S3).

### Composition of CHD4-NURD complexes during spermatogenesis

NURD function is influenced by its subunit composition [17, 18]. To determine whether the expression of NURD composition might change during spermatogenesis, we analyzed the levels of representative NURD subunits in enriched fractions of undifferentiated and differentiating spermatogonia, as well as in the flow-through (enriched in Sertoli cells) after spermatogonia enrichment. NURD subunits HDAC2A, MTA1, RBBP4, RBBP7 and MBD2 were present in enriched fractions of undifferentiated (THY1.2+) and differentiating (c-KIT+) spermatogonia (Fig. 1B), suggesting a particular composition and perhaps a different function for CHD4-NURD in this cell type.

To determine the composition of CHD4-NURD complexes, we used co-immunoprecipitation analysis to uncover NURD subunits that interact with CHD4 in enriched fractions of THY1.2 and c-KIT spermatogonia cells together and the flowthrough. The NURD subunits HDAC2A, MTA1, and RBBP4/RBBP7, but not MBD2, coimmunoprecipitated with CHD4 from all fractions (Fig. 1C). HDAC2A coimmunoprecipitated with CHD4 from wild type and *Chd3^-/-^* spermatogonia cells (Fig. S2B). These data suggest that: i) CHD4 forms a NURD complex independently of CHD3, ii) that loss of CHD3 does not perturb CHD4-NURD complex formation in spermatogonia cells.

### Deletion of *Chd4* but not *Chd3* results in testis developmental defects

To examine the potential functions of *Chd4* and *Chd3* during spermatogenesis, we generated a series of *Chd4* and *Chd3* germline conditional knockout mice (Fig. 2). To delete the floxed allele in gonocytes (embryonic day 15.5, Fig. 2C) [19], male Ddx4-Cre; *Chd4^WT/Δ^* were crossed with *Chd4^fl/fl^* females to generate Ddx4-Cre; *Chd4^fl/Δ^* conditional knockout mice (here called *Ddx4-Chd4^-/-^*) (Fig. 2A). A similar strategy was used to generate Ddx4-*Chd3^-/-^* mice (Fig. 2B).

**Figure 2.**
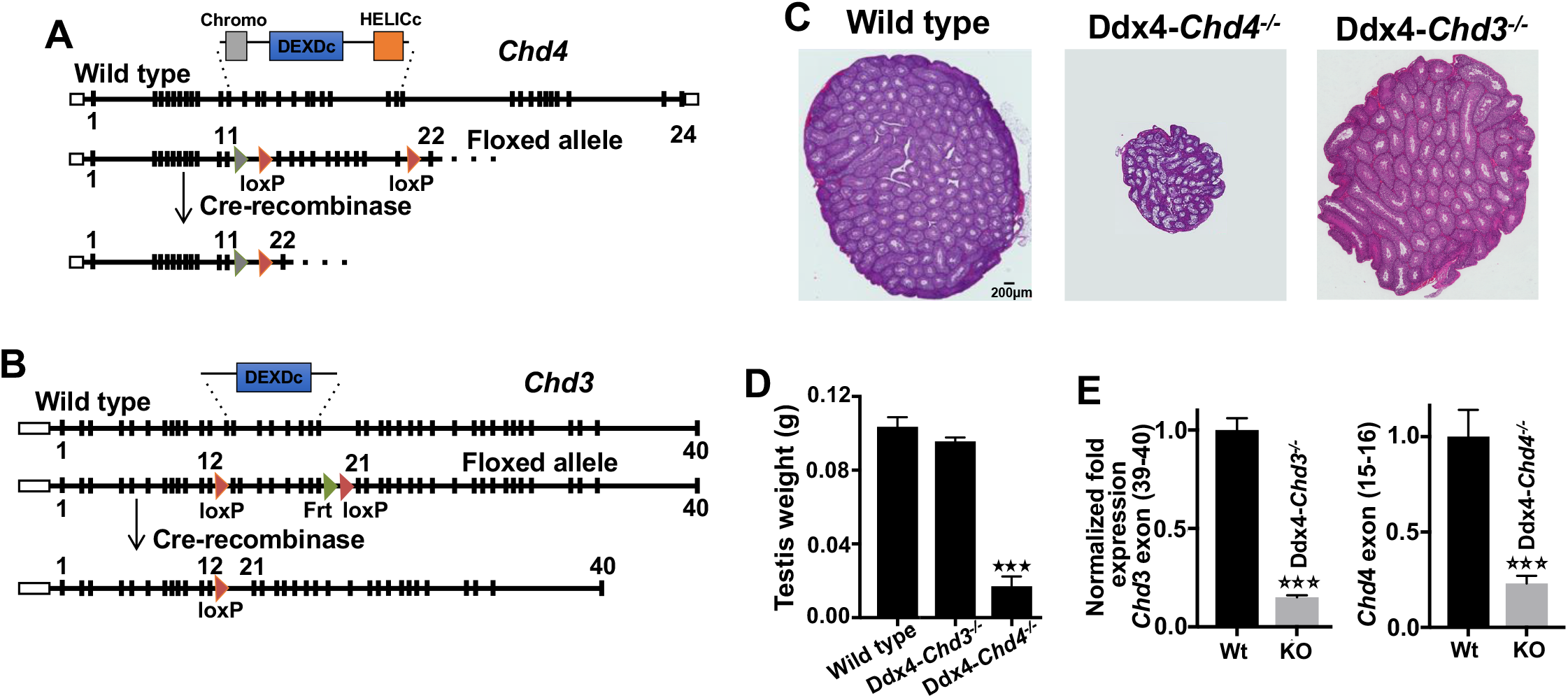
*Chd4* and *Chd3* gene targeting design and testis developmental defects in *Chd4* mutant mice. **(A and B)** Testis specific Cre knockout strategy for deletion of *Chd4* and *Chd3*. See description in Materials and Methods. **(C)** H&E-stained paraffin testis sections of wild type, Ddx4-*Chd4^-/-^* and Ddx4-*Chd3^-/-^* mice. **(D)** Quantification of testis weight for wild type and homozygous knockout mice is shown. Images and testis weigh measurements were obtained from three mice. **(E)** *Chd3* transcription levels were measured in Ddx4-*Chd3^-/-^* total testis and compared to wild type littermates (two months old mice, n=3). Chd4 transcription level in enriched fractions of spermatogonia obtained from 7 dpp *Ddx4-Chd4^-/-^* and control wild type litter mate mice (mice, biological replicate n=3).

*Ddx4-Chd4^-/-^* adult mice (2 months old) appeared normal in all aspects except in the reproductive tissues. Testes were significantly smaller in Ddx4-*Chd4^-/-^* males (mean: 0.017g ± SD: 0.005, number of quantified mice n□=□4 (8 testes), P≤0.0001, t test) compared to wild type (0.104 g ± 0.015, n□=□6 mice (12 testes)) littermates (Fig. 2C and D), indicating severe developmental defects in the testis. We confirmed deletion of *Chd3* and *Chd4* by RT-qPCR (Fig. 2E). Immunofluorescence analysis of neonatal Ddx4-*Chd4^-/-^* testis CHD4 was absent at 1ddp (Fig. S2A).

We found that adult Ddx4-*Chd4^-/-^* males develop testicular hypoplasia with hyperplasia of interstitial cells and lack spermatozoa (Fig. 3A). The number of seminiferous tubules is similar between wild type and mutant animals, but the diameter is reduced (wild type, mean ± SD, 287 ± 34, n=400 seminiferous tubules cross sections (3 different mice, 2-month-old) versus *Chd4^-/-^* 148 ± 21.3, n=210, P<0.0001 t test).

**Figure 3.**
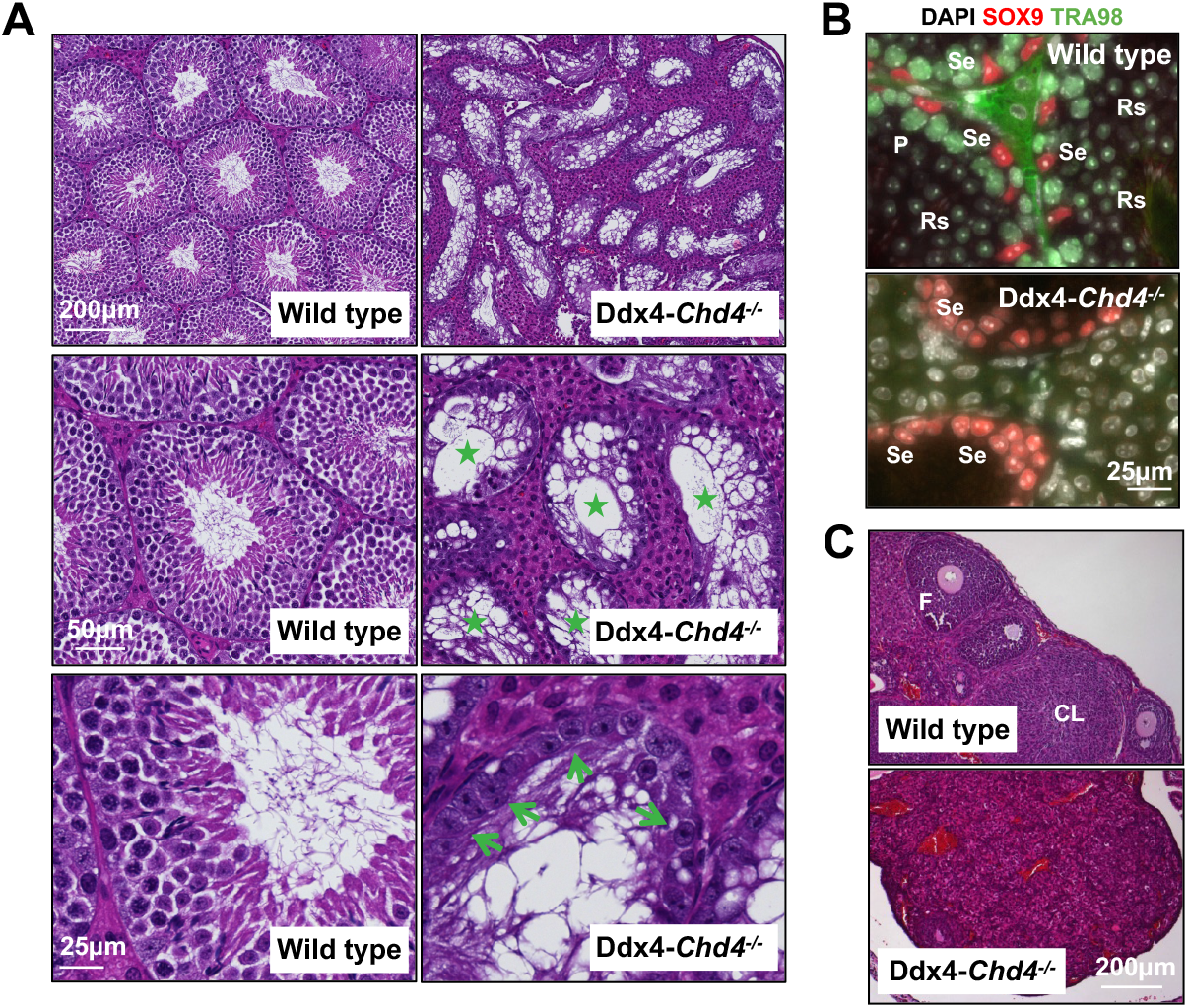
*Ddx4-Chd4^-/-^* mice show profound defects in gametogenesis. **(A)** H&E-stained histological sections of wild type and Ddx4-*Chd4^-/-^* testis. Stars mark seminiferous tubules with absent germ cells. Note unchanged number and morphology of Sertoli cells (indicated by green arrows). **(B)** Histological sections of wild type and *Ddx4-Chd4^-/-^* testis showing seminiferous tubules immunolabeled with SOX9 (a marker of Sertoli cells) and TRA98 (to mark germ cells). P, pachytene cells; Se, Sertoli cells; Rs, rounded spermatids. **(C)** H&E-stained histological sections of wild type and Ddx4-*Chd4^-/-^* ovaries. F, follicles and CL, corpora lutea.

Analysis of Ddx4-*Chd4^-/-^* testes revealed a total loss of germ cells (marked by TRA98) in seminiferous tubules (Fig. 3B). No developing gametes were observed, including cell types at early stages (e.g., spermatogonia) (Fig. 3A and B). Sertoli cells develop normally in Ddx4-*Chd4^-/-^* mice, consistent with the specific loss of *Chd4* in germ cells at early stages of development. We did not observe differences in germ cell development between wild type and Ddx4-*Chd3^-/-^* mice (Fig. S3), consistent with their similar testis sizes (Fig. 2C and D).

We also analyzed H&E-stained histological sections of ovaries from 45-day-old wild type and Ddx4-*Chd4^-/-^* female mice. We noted a significant reduction in ovary size, an increase in stromal cells, a reduced number of follicles (wild type, 10 ± 3, n=6 mice versus *Ddx4-Chd4^-/-^* 0.6 ± 0.9, n=5, P<0.0001 t test) and absent corpora lutea (wild type, 9 ± 1, n=6 mice versus Ddx4-*Chd4^-/-^* 0 ± 0, n=5, P<0.0001 t test) in the Ddx4-*Chd4^-/-^* mice compared to wild type (Fig. 3C).

We conclude that deletion of *Chd3* has no apparent effect on gamete development. However, germ cell specific deletion of *Chd4* results in severe male and female germ cell developmental defects, possibly originated at premeiotic stages of development.

### CHD4 is required for neonate spermatogonia survival

The severe phenotypes observed in Ddx4-*Chd4^-/-^* mice (Fig. 2 and 3) prompted us to investigate spermatogonial differentiation during testis development in newborns. Testis sections from 9 dpp Ddx4-*Chd4^-/-^* mice stained with H&E showed a markedly reduced number of germ cells (Fig. 4A) as well as differences in cell composition, compared to those from age-matched wild type mice. To analyze this in detail, we examined the presence of cells expressing STRA8 (Fig. 4A), which marks differentiating spermatogonia, SYCP3 and yH2AX which are markers of primary spermatocytes and TRA98, a marker for germ cells (Fig. S5). Whereas tubules from 9 dpp wild type mice contained cells expressing TRA98 (45 ± 10, n=66 seminiferous tubules counted obtained from 3 mice) and STRA8 (18.8 ± 8.4, n=36 obtained from 3 mice), tubules from *Chd4^-/-^* mice showed a near absence of cells expressing these markers (TRA98 4.6 ± 3, n=60 obtained from 3 mice, P<0.0001, t test; STRA8 2.5 ± 3.5, n=42 obtained from 3 mice, P<0.0001, Student t test) (Fig. 4A). Testes sections from 9 dpp *Chd4^-/-^* mice also showed a reduction in primary spermatocytes expressing the meiotic prophase I markers SYCP3 and γH2AX compared to those from 9 dpp wild type mice (Fig. S5A and B). Together, the results further suggest that testis defects in *Chd4^-/-^* mice begin early, during pre-meiotic stages of postnatal development, leading to an absence of germ cells in adults.

**Figure 4.**
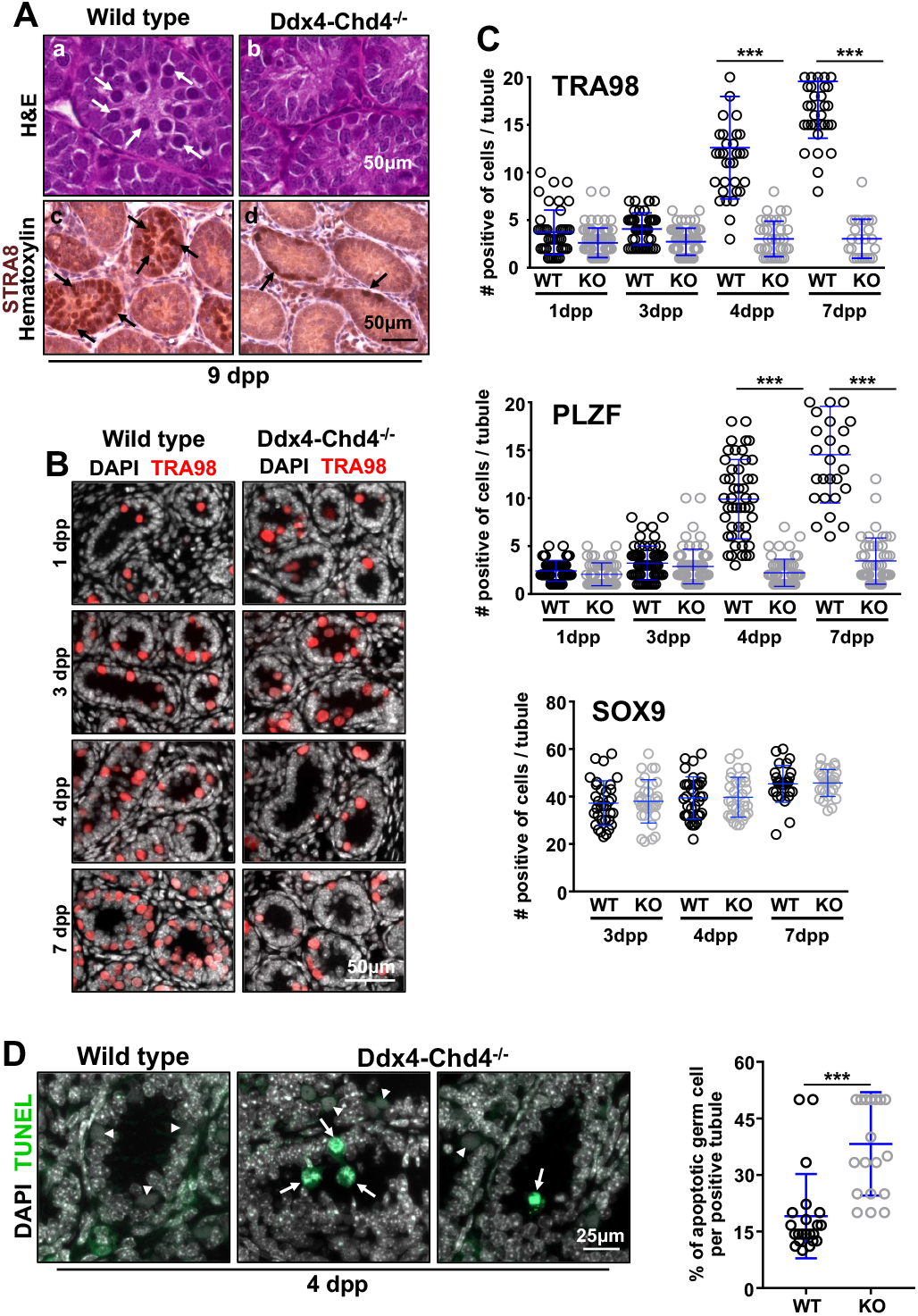
Spermatogonia cell survival requires CHD4. **(A)** Histological sections of wild type and Ddx4-*Chd4^-/-^* testis cord from 9 days old mice stained with H&E (a and b) and Hematoxylin and immunostained with STRA8 antibodies (marking differentiating spermatogonia) (c and d). Arrows mark type of cells present in wild type but reduced in number or absent in the mutant. Experiments were done in at least three different mice showing similar results. **(B)** Immunostaining of sections of developing testis reveals severe loss of germ cells (TRA98) in *Chd4^-/-^* testis cords. **(C)** Quantitation of number of cell positive for TRA98, PLZF, and SOX9. 4dpp testis (PLZF, wild type, 9.9 ± 4.1, n=47 seminiferous tubules; *Chd4^-/-^* 2.2 ± 1.4, n=56, P<0.0001, t test. TRA98, wild type 12.6 ± 5.3, n=35; *Chd4^-/-^ 3.0* ± 1.9, n=37, P<0.0001, t test). 7dpp testes (PLZF, wild type, 14.6 ± 5.0, n=28; *Chd4^-/-^* 3.4 ± 2.4, n=45, P<0.0001, t test. TRA98, wild type 19.6 ± 5.9, n=48; *Chd4^-/-^* 3.0 ± 2.1, n=24, P<0.0001, t test). Experiments were done in three different mice showing similar results combined in our quantification. **(D)** Higher cell death in *Chd4^-/-^* knockout testes than in wild type controls at four days post-partum. Apoptotic cells were visualized by staining for TUNEL in three mice of each genotype. The percentage of apoptotic cells was counted only in the tubule cross-sections that contained apoptotic cells and was normalized to total number of germ cells. 19.05 ± 11.17, n = 468 (mean ± SD, n= number of seminiferous tubules counted) wild type and 38.25 ± 13.72 *Chd4^-/-^*, n = 344; P<0.0001 (two-tailed Student t test). The arrows indicate apoptotic cells and arrowhead non-apoptotic germ cells.

To pinpoint when the testes defects originate in *Ddx4-Chd4^-/-^* mice, we stained testes sections from 1-21 dpp mice for the expression of TRA98 (all germ cells). We observe that as general trend, in all analyzed stages, the number of TRA98 positive cells was reduced in *Chd4^-/-^* mice compared to wild type, progressing to total absence of germ cells (Fig. S6). *Chd4* mutant sections displayed a small reduction in the number of cells expressing TRA98 at both 1dpp and 3dpp (Fig. 4B and 4C and Fig. S6). We observed equal numbers of SOX9-positive Sertoli cells in testes from wild type and *Chd4^-/-^* mice at 3, 4, and 7 dpp (Fig. 4C), as expected for the specific loss of *Chd4* in spermatogonia cells. Both PLZF-positive and TRA98-positive cells were substantially reduced in *Chd4^-/-^* testis compared to wild type testis at 4 dpp and 7dpp (Fig. 4C).

Given that Chd4 may act as a regulator of cell-cycle progression, we then examined whether the rapid loss of PLZF-positive neonate spermatogonia in *Chd4^-/-^* testes was due to altered proliferative activity. We conducted EdU incorporation study to test this possibility. 4 dpp mice were injected with EdU and analyzed 3 h later, after which we assayed its incorporation in PLZF-positive spermatogonia in whole mounts of seminiferous tubules (Fig. S7A). We found that spermatogonia cell proliferation (PLZF/EdU^+^ cells) in wild type and Ddx4-Chd4^-/-^ is proportionally the same (Fig. S7C). In addition, reduced amount of total PLZF-positive cells was found in Ddx4-CHD4^-/-^ compared to wild type in the whole-mounting experiment (Fig. S7B).

To determine whether cell death contributed to the loss of Ddx4-*Chd4^-/-^* neonate spermatogonia (Fig. 4C), we performed TUNEL assay in staining paraffin embedded testis sections of wild type and *Chd4^-/-^* four days old testis. At this age the testis is mostly constituted by spermatogonia and Sertoli cells, which can be easily distinguished by DAPI nuclear staining patterns. We found a significant increase in the percentage of apoptotic cells in *Chd4^-/-^* testis compared to wild type mice (Fig. 4D).

We conclude that the possible cause of spermatogonia failure in *Chd4^-/-^* mice is in the survival/maintenance of neonate undifferentiated spermatogonia.

### CHD4, DMRT1, and PLZF work together in a regulatory axis involved in spermatogonia cell survival

Our results show that CHD4 is required for spermatogonia maintenance/survival. We then reason that CHD4 may interact with genes that have been described to work in spermatogonia maintenance. Indeed, DMRT1 has been show to function in spermatogonia stem cells maintenance, and this function seems to be mediated by direct regulation of Plzf gene expression, another transcription factor required for spermatogonia maintenance [20]. To test our hypothesis, we immunostained paraffined testes sections from 1, 4, and 7 dpp mice for the presence of DMRT1 (Fig. 5A and B). Compared to wild type, *Chd4* mutant showed a clear reduction in DMRT1 immunosignal.

**Figure 5.**
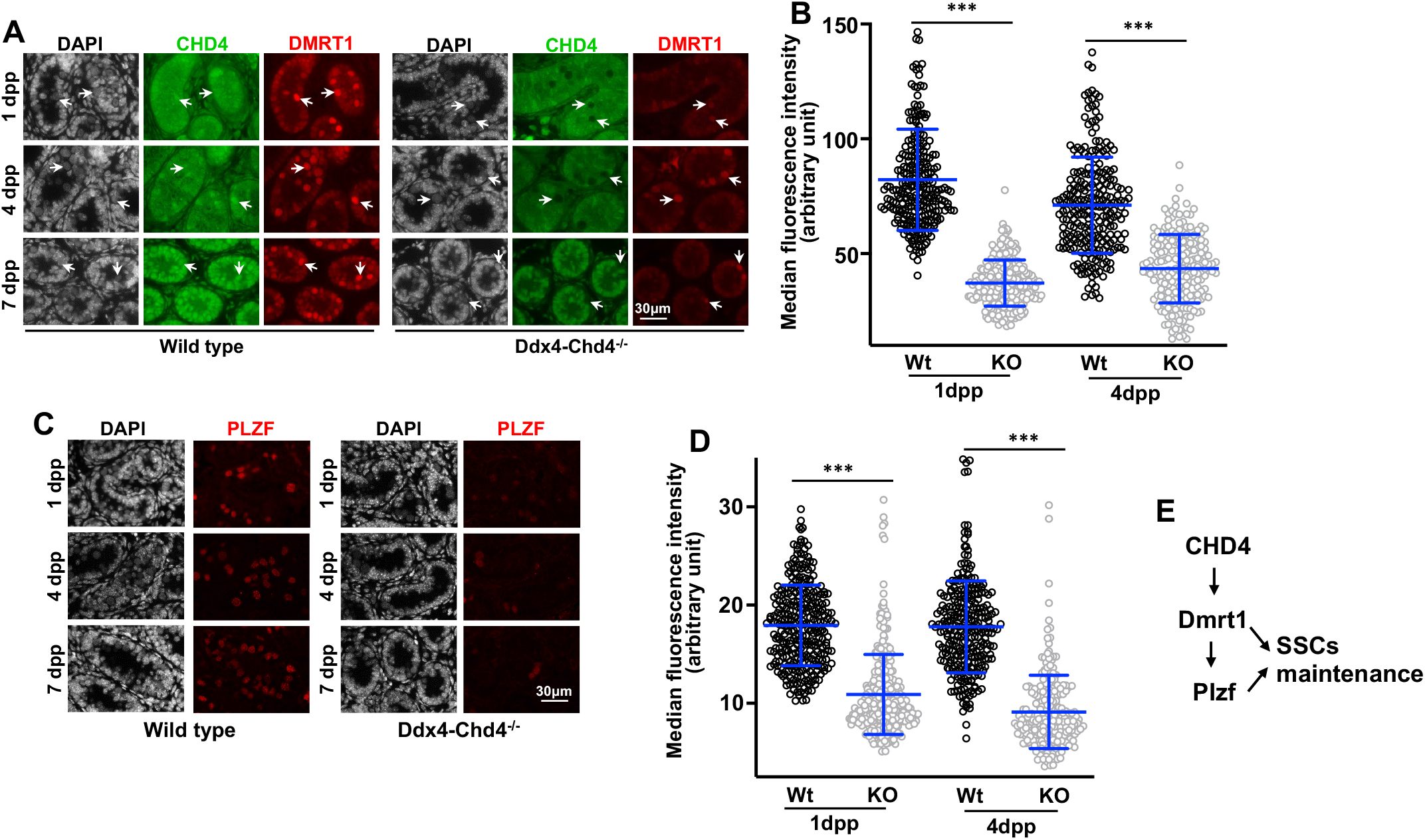
Effect of Chd4 deletion in Dmrt1 and Plzf expression. **(A)** Histological sections of wild type and Ddx4-*Chd4^-/-^* testis from 1, 4, and 7 dpp mice stained with CHD4 and DMRT1 antibodies. Arrows indicate examples of CHD4 positive spermatogonia (Wt) or CHD4 absence of staining in Ddx4-*Chd4^-/-^* spermatogonia and the correspondent cell stained for DMRT1. **(B)** Quantitation of fluorescence intensity of DMRT1 expression on spermatogonia is shown for wild type (Wt) and Ddx4-*Chd4^-/-^* knockout at ages of 1 dpp (Wt = 82.17 ± 22.02, n=272; and KO = 37.20 ± 10.01, n=287, P<0.0001, Student t test) and 4 dpp (Wt = 71.09 ± 20.93, n=242; and KO = 43.47 ± 14.88, n=198, P<0.0001, Student t test). Values represent the median fluorescence intensity ± standard deviation, n = total number of cells analyzed from 3 biological replicates using 3 different mice. (**C**) Histological sections of wild type and Ddx4-*Chd4^-/-^* testis from 1, 4, and 7 dpp mice stained with antibodies against PLZF. **(D)** Quantitation of fluorescence intensity of PLZF expression on spermatogonia is shown for wild type and Ddx4-*Chd4^-/-^* knockout at ages of 1 dpp (wt = 17.94 ± 4.10, n=300; and Ddx4-*Chd4^-/-^* = 10.89 ± 4.07, n=339, P<0.0001, Student t test) and 4 dpp (Wt = 17.78 ± 4.70, n=282; and Ddx4-*Chd4^-/-^* = 9.11 ± 3.75, n=230, P<0.0001, Student t test). Values represent the median fluorescence intensity ± standard deviation, n = total number of cells analyzed from 3 biological replicates using 3 different mice. **(E)** Diagram representing CHD4, Dmrt1, and Plzf relationship in spermatogonia stem cell maintenance. Note that our studies do not address the possibility that Plzf may be a direct target of CHD4.

Recent work described that Dmrt1 controls Plzf expression, which is a transcription factor required for spermatogonia maintenance [20]. We then tested the effect of CHD4 depletion in Plzf expression. PLZF immunostaining of testes sections from 1, 4, and 7 dpp mice revealed a significantly reduction of this protein in Chd4 knockout cells respect to wild type (Fig. 5C and D). This is consistent with a model by which CHD4 control of Dmrt1 and downstream targets influences cell survival/maintenance (Fig. 5E).

We concluded that Chd4 participates in the maintenance/survival of neonate spermatogonia stem cell possibly through transcriptional regulation of genes participating in these critical processes. We note, however, that the dramatic phenotype observed in CHD4^-/-^ spermatogonia likely reflect CHD4 targeting a wide spectrum of genes participating in different pathways.

## Discussion

In this work, we examined the potential function of two critical NURD catalytic subunits, CHD4 and CHD3, in spermatogenesis. Our data suggest that a CHD4-NURD (but not CHD3-NURD) complex controls neonate spermatogonia development at early stages of testis development. Germline deletion of *Chd4*, but not *Chd3*, results in a severe loss of germ cells specifically at early stages of the testis cord development. *Chd4* deletion affects spermatogonia, with the first obvious consequences in undifferentiated spermatogonia.

Most CHDs are expressed in testis; however, their insertion into the NURD complex seems to be developmentally regulated, with apparent different patterns of expression during gametogenesis. CHD5 [7–9] has been shown to be expressed and required to compact chromatin in postmeiotic stages of spermatogenesis [12, 21]. Our results show that CHD4 is expressed and functions at premeiotic stages of gametogenesis.

The functions of CHD4-NURD in neonate spermatogonia development and male gametogenesis are further revealed by our cytological analysis showing that CHD4 regulates the expression of Dmrt1 (Fig. 5). DMRT1 function in maintenance of spermatogonia stem cells has been proposed to be mediated by direct regulation of Plzf gene expression, another transcription factor required for spermatogonia maintenance [20]. In agreement with this possibility, we observed that the amount of PLZF was significantly reduced in Chd4 knockout versus wild type cells. In sum, our work provides evidence of a regulatory axis in which CHD4 may control important genes involved in spermatogonia stem cell maintenance and survival. Additional work will be required to test CHD4 effect in Sohlh1, another Dmrt1 direct target involved in cell survival and differentiation [20, 22, 23], and the Stra8 gene, the latter which precocious expression is detected in Dmrt1 knockout mice. We note that the dramatic cellular phenotype we observed after CHD4 depletion in spermatogonia may be only explained by CHD4 activity targeting several genes and different pathways.

## Material and Methods

### Mice

*CHD4-floxed* mice (*CHD4^fl/fl^*) have been described [24]. *Chd3-*floxed mice were generated by Cyagen Biosciences using homologous recombination of a targeting vector in C57Bl/6 embryonic stem cells. The targeting vector incorporated a 5’ LoxP site inserted between exons 12 and 13 and a 3’ LoxP site inserted between exons 20 and 21 of the wild type *Chd3* allele. Transgenic *Cre* recombinase mice *Ddx4-Cre*^FVB-Tg(Dd×4-cre)1Dcas/J^ was purchased from The Jackson Laboratory (Bar Harbor, ME). *Chd3 or Chd4* gonad-specific knockouts and wild-type heterozygotes littermates were obtained from crosses between female homozygous *flox/flox* mice with male heterozygous Cre/+; *Chd3* Wt/flox and/or *Chd4* Wt/flox mice.

All experiments conformed to relevant regulatory standards guidelines and were approved by the Oklahoma Medical Research Foundation-IACUC (Institutional Animal Care and Use Committee).

### Mice Genotyping

Characterization of wild type and *floxed* alleles was carried out by PCR using the following oligonucletides: CHD3 forward 5’-GGGTGGAGGTGGAAAGTGTA, CHD3 reverse 5’-AGAGGACAGGTCACAGGACAA, CHD4 forward 5’-TCCAGAAGAAGACGGCAGAT and CHD4 reverse 5’-CTGGTCATAGGGCAGGTCTC. The presence of *cre* recombinase allele were determine by PCR using the following primers: DDX4-Cre forward 5’-CACGTGCAGCCGTTTAAGCCGCGT, DDX4-Cre reverse 5’-TTCCCATTCTAAACAACACCCTGAA.

### Real time-PCR

Total RNA was isolated from adult testis or from enriched fractions of spermatogonia with the Direct-zol RNA MiniPrep Plus kit (Zymo Research). RNA (2.0μg) was oligo-dT primed and reverse-transcribed with the high-capacity RNA-to-cDNA kit (Applied Biosystems). Exon boundaries of *Chd4 and Chd3* were amplified using TaqMan Assays (Applied Biosystems) as directed by the manufacturer using Beta-2 macroglobulin as standard. TaqMan Mm01190896_m1 (Chd4), Mm01332658_m1 (Chd3), and Mm00437762_m1 (Beta-2 microglobulin). Gene expression was normalized with respect to wild type with wild type expression levels considered to be 1.

### Western blot cell lysates

Total testis or enriched cells fractions were lysed in ice-cold protein extraction buffer containing 0.1% Nonidet P-40, 50 mM Tris-HCl, pH 7.9, 150 mM NaCl, 3 mM MgCl2, 3 mM EDTA, 10% glycerol, 1 mM DTT, 1mM PMSF and protease inhibitors (ThermoFisher Scientific, A32965) followed by sonication (3 pulses of 10 seconds) using micro ultrasonic cell disrupter (Kontes). The relative amount of protein was determined measuring absorbance at 260nm using NanoDrop 2000c spectrophotometer (ThermoFisher Scientific). Proteins were solubilized with 2X sample buffer (4% SDS, 160 mM Tris-HCl, pH 6.8, 20% glycerol, 4% mM β-mercaptoethanol, and 0.005% bromophenol blue) and 30 μg/lane of sample were separated by 4–15% gradient Trisacetate SDS-PAGE and electro transferred to PVDF membrane (Santa Cruz Biotechnology, sc-3723). The blots were probed with individual primary antibodies, and then incubated with HRP-conjugated goat anti-mouse or rabbit antibody as required. In all blots, proteins were visualized by enhanced chemiluminescence, and images acquired using Western Blot Imaging System c600 (Azure Biosystems). ImageJ software were used for quantification of non-saturated bands and α-tubulin were used for normalization. Antibodies used are detailed in table S1.

### Histology and immunostaining

Testes and ovaries were dissected, fixed in 10% neutral-buffered formalin (Sigma) and processed for paraffin embedding. After sectioning (5–8-μm), tissues were positioned on microscope slides and analyzed using hematoxylin and eosin using standard protocols. For immunostaining analysis, tissue sections were deparaffinized, rehydrated and antigen was recovered in sodium citrate buffer (10 mM Sodium citrate, 0.05% Tween 20, pH 6.0) by heat/pressure-induced epitope retrieval. Incubations with primary antibodies were carried out for 12 h at 4°C in PBS/BSA 3%. Primary antibodies used in this study are detailed in table S6 Following three washes in 1 X PBS, slides were incubated for 1 h at room temperature with secondary antibodies. A combination of fluorescein isothiocyanate (FITC)-conjugated goat anti-rabbit IgG (Jackson laboratories) with Rhodamine-conjugated goat anti-mouse IgG and Cy5-conjugated goat anti-human IgG each diluted 1:450 were used for simultaneous triple immunolabeling. Slides were subsequently counterstained for 3 min with 2 μg/ml DAPI containing Vectashield mounting solution (Vector Laboratories) and sealed with nail varnish. We use Zen Blue (Carl Zeiss, inc.) for imaging acquisition and processing.

### Enrichment of Spermatogonia Populations

Our procedure of cell enrichment followed [25, 26]. Briefly, testis from 7 dpp mice (or any other indicated age) were removed from mice and placed in a Petri dish containing Dulbecco’s Modified Eagle Medium (DMEM without phenol red). After detachment of the tunica albuginea, the seminiferous tubules were loosen using forceps and incubated in a 15 ml tube containing DMEM containing 1 mg/mL of collagenase, 300 U/mL of hyaluronidase and 5 mg/mL DNAse I (StemCell Technologies) under gentile agitation for 10 min. The seminiferous tubule clumps were pelleted by gravity and the cell suspension containing interstitial cells was discarded. The tubules were then incubated of with 0.05% Trypsin-EDTA solution (Mediatech Inc) for 5 min and the reaction was stopped by adding 10% volume of 10% BSA in PBS. Single cell suspension was obtained by mechanical resuspension followed by filtration through a 40-μm-pore-size cell strainer and dead cel;ls were removed using Dead Cell Removal Kit (Miltenyi Biotec 130-090-101). Differentiating c-KIT+ neonate spermatogonia cells were magnetically labeled with CD117 (c-KIT+) MicroBeads (Miltenyi Biotec 130-091-224) and isolated using MS columns (Miltenyi Biotec 130-042-201) according to manufacturer’s instructions. After the depletion of the c-KIT+ cells, the population of undifferentiated neonate spermatogonia cells were separated using CD90.2 (THY1.2+) MicroBeads (Miltenyi Biotec 130-121-278). Relative enrichment of cell populations was evaluated by STRA8 (c-Kit fractions) or PLZF (THY1.2 fractions) western blots (Fig. 1B). After c-kit and THY1.2 separation, the flow-through mostly contained Sertoli cells (SOX9 positive, Fig. S3). The number of cells obtained from a pool of 4 mice testis at 7dpp was approximately 3.43×^5^ in THY1.2 fractions and 5.71×10^5^ in c-Kit fractions.

### Immunoprecipitation

Co-immunoprecipitation experiments were performed using testis of wild type or Ddx4-CHD3^-/-^ mouse (Adult - 2 months old). After detunication, seminiferous tubules were loosen using forceps, washed twice with cold PBS and lysed using ice-cold RIPA buffer (50 mM Tris pH 7.5, 150 mM NaCl, 1 % NP40, 0.5 % Deoxycholate) containing protease inhibitors (ThermoFisher Scientific, A32965), sheared using 23 G needle, incubated on ice for 15 min and centrifugated at 1000 x g for 10 minutes at 4 °C. Supernatant were collected in a separate tube, the pellet were resuspended in RIPA buffer, disrupted by sonication (3 pulses of 10 seconds) and centrifuged 12.000 x g. This second supernatant was combined with the previous one and protein concentration was determined. We used 1mg of protein for each immunoprecipitation. Lysates were pre-cleared with protein G magnetic beads (BioRad, 161-4023) for 1 hour at room termperature and incubated with rabbit anti-CHD3 (5μg, Bethyl A301-220A), rabbit anti-CHD4 (2 μg, Abcam ab72418), or rabbit IgG (5μg Jackson ImmunoResearch, 011-000-003). Lysates were rotated overnight at 4 °C and immune complexes were collected with protein G magnetic beads (2 hours at 4 °C). Beads were washed 4 times with washing buffer (50 mM Tris pH 7.5, 150 mM NaCl, 0.1 % TX100, 5 % glycerol) and two times with PBS. Proteins were eluted by boiling the beads with 2X sample buffer and analyzed by SDS-PAGE as described above.

### EdU-based proliferation assay

Mice at indicated age received subcutaneous injection of EdU (50 mg/Kg) (Invitrogen, A10044) 3 hours prior euthanasia. After that, testes were removed and processed for whole-mount immunohistochemistry. EdU was detected by incubation of testis samples with reaction mix (2 mM CuSO_4_, 50 mM ascorbic acid and 2 mM Alexa Azide conjugates (488 or 647) in PBS) for 3 hours at room temperature.

### Whole-mount seminiferous tubules

Immunohistochemistry of whole-mount seminiferous tubules was performed as described [27]. Briefly, after detachment of the tunica albuginea, the seminiferous tubules were loosen using forceps and incubated in a 15 ml tube containing DMEM containing 1 mg/mL of collagenase, 300 U/mL of hyaluronidase and 5 mg/mL DNAse I (StemCell Technologies) under gentile agitation for 10 min. The seminiferous tubules clumps were pelleted by gravity and the cell suspension containing interstitial cells was discarded. Seminiferous tubules were fixed for 4h in 4 % PFA (pH7.2 in PBS) at 4 °C. After extensively wash in PBS, the tubules were permeabilized with series of MeOH/PBS (25, 50, 75, 95 %, and twice in 100 % MeOH) for 15 min at room temperature, treated with MeOH: DMSO: H_2_O_2_ (4:1:1), and rehydrated with MeOH/PBS (50, 25 % and twice in PBS). Samples were incubated in ice-cold blocking solution PBSMT (PBS with 2 % non-fat dry milk and 0.5 % triton X-100) for 3 hours and then over-night at 4 °C with indicated primary antibodies under gentle rotation. Seminiferous tubules were washed in PBSMT (5 x 1h) and incubated with dye conjugated (Alexa488 or TRITC) goat antimouse or rabbit antibody as required. The tubules were mounted in raised coverslips glass slides.

### Statistical Analyses

Results are presented as mean ± standard deviation (SD). Statistical analysis was performed using Prism Graph statistical software. Two-tailed unpaired Student’s t-test was used for comparisons between 2 groups. P < 0.05 was considered statistically significant.

