## supplementary legend for "Mouse CHD4-NURD is required for neonatal spermatogonia survival and normal gonad development"

**Supplementary Information**

**Supplementary Figures**

**Figure S1. CHD4 expression during gametogenesis.** Expression of *Chd4* monitored by immunofluorescence in paraffin embedded testis sections of 1, 3, 4, 7, 9, 14, and 120 dpp mice. Spermatogonia cells are positive for PLZF and Sertoli cells are positive for SOX9.

**Figure S2. Specificity of CHD4 immunostaining and formation of CHD4- and CHD3-NURD complexes in developing gametes. (A)** Antibodies against CHD4 colocalize with PLZF positive cells in seminiferous tubules of 1 dpp wild type mice but no signal of CHD4 is detected in Ddx4-*Chd4^-/-^* mice. Note Sertoli cells (SOX9) exhibit CHD4 immunosignal in both wild type and Ddx4-*Chd4^-/-^* (Ddx4) mice. This is a representative image out of three different experiments using one wild type or Ddx4-*Chd4^-/-^* per experiment. Note that the red channel (PLZF staining) in the image corresponding to Ddx4-*Chd4^-/^*^-^ was exposed longer to allow comparison to wild type. **(B)** CHD4 co-immunprecipitates HDAC2A, a core component of the NURD complex, independent of CHD3. The asterisk indicates unspecific signal in the CHD3 blot and arrow indicates the specific band.

**Figure S3. Spermatogonia enrichment. Analysis of THY1+ and c-KIT+ fractions**. Cells were attached to coverslips and immunostained with the indicated antibodies. **(A)** Quantification of germ cell shows that spermatogonia represents 61.5% (±3.7%) from THY1+ fractions (PLZF positive cells) and 65.4% (±11%) from c-KIT fractions (DDX4 positive cells). **(B)** The flow-through fraction obtained after use of the THY1 and c-KIT columns is composed of 60.6% ± 3.6% Sertoli cells. A minor fraction of Sertoli cells was also detected in THY1+ (28.1 ± 6.3%) and c-KIT+ (25.4% ± 7.4%) fractions. Numbers represent average ± standard deviation from 2 biological replicates and 4 technical replicates.

**Figure S4.** **CHD3 is dispensable for gametogenesis.** H&E stained histological sections of wild type and Ddx4-*Chd3^-/^*^-^ testis. No differences in type or number of germ cell at any stage of development are observed between wild type and Ddx4-*Chd3^-/^*^-^ knockouts.

**Figure S5. *Chd4* deletion results in deficient spermatogonia cell development and near absence of spermatocytes. (A)** Histological sections of 9 dpp wild type and Ddx4-*Chd4^-/^*^-^ testis showing seminiferous tubules immunolabeled with TRA98 (a marker of germ cells) and SYCP3 and γH2AX (markers of primary spermatocytes). Arrows indicate examples of positive cells. **(B)** Quantification of cell number per positive tubule shown in A. TRA98 positive cells in wild type (45 ± 10, n=66) and Ddx4-*Chd4^-/^*^-^ (4.6 ± 3, n=60, P<0.0001, Student t test) mice. SYCP3 positive cells in wild type (25 ± 5, n=45) and Ddx4-*Chd4^-/^*^-^ (2 ± 1, n=45, P<0.0001, Student t test) mice. γH2AX positive cells in wild type (20 ± 7, n=38) and Ddx4-*Chd4^-/^*^-^ (2 ± 1, n=38, P<0.0001, Student t test) mice.

**Figure S6. Deficient spermatogonia cell survival and differentiation in *Chd4^−/−^*** **mice.** Histological sections of wild type and Ddx4-*Chd4^-/^*^-^ testis cords from 1, 3, 4, 7, 9, 14, and 21 dpp mice stained with TRA98 antibodies. See quantification in Figure 4C.

**Figure S7. Deficient spermatogonia cell survival in *Chd4^-/^*^-^** **knockout mice.** **(A)** Immunostaining of whole mount seminiferous tubules reveals loss of spermatogonia (PLZF) in 4dpp DDX4-*Chd4^-/^*^-^ mice. EdU was used to mark proliferating cells. **(B)** Quantitation of number of PLZF+ cells per mm of seminiferous tubules (wild type (116 ± 48) and Ddx4-*Chd4^-/^*^-^ (54 ± 22), P<0.0011, student t test). Number of cells corrected by total length of seminiferous tubule analyzed in wild type versus *Chd4^-/^*^-^ mutants. A total of 7.36 mm (833 cells, wild type) and 10.15 mm (478 cells, DDX4-*Chd4^-/^*^-^) seminiferous tubule length was counted using four different mice. **(C)** Percentage of proliferative spermatogonia cells (PLZF^+^/EdU^+^) in wild type and DDX4-*Chd4^-/^*^-^ (wild type (63% ± 13%) and DDX4-*Chd4^-/^*^-^ (56% ± 16), P<0.2816, Student t test).

**Table S1. List of antibodies used in this work.**
