## Supplementary figures and images for "Mouse CHD4-NURD is required for neonatal spermatogonia survival and normal gonad development"

### Supplementary Figure 1

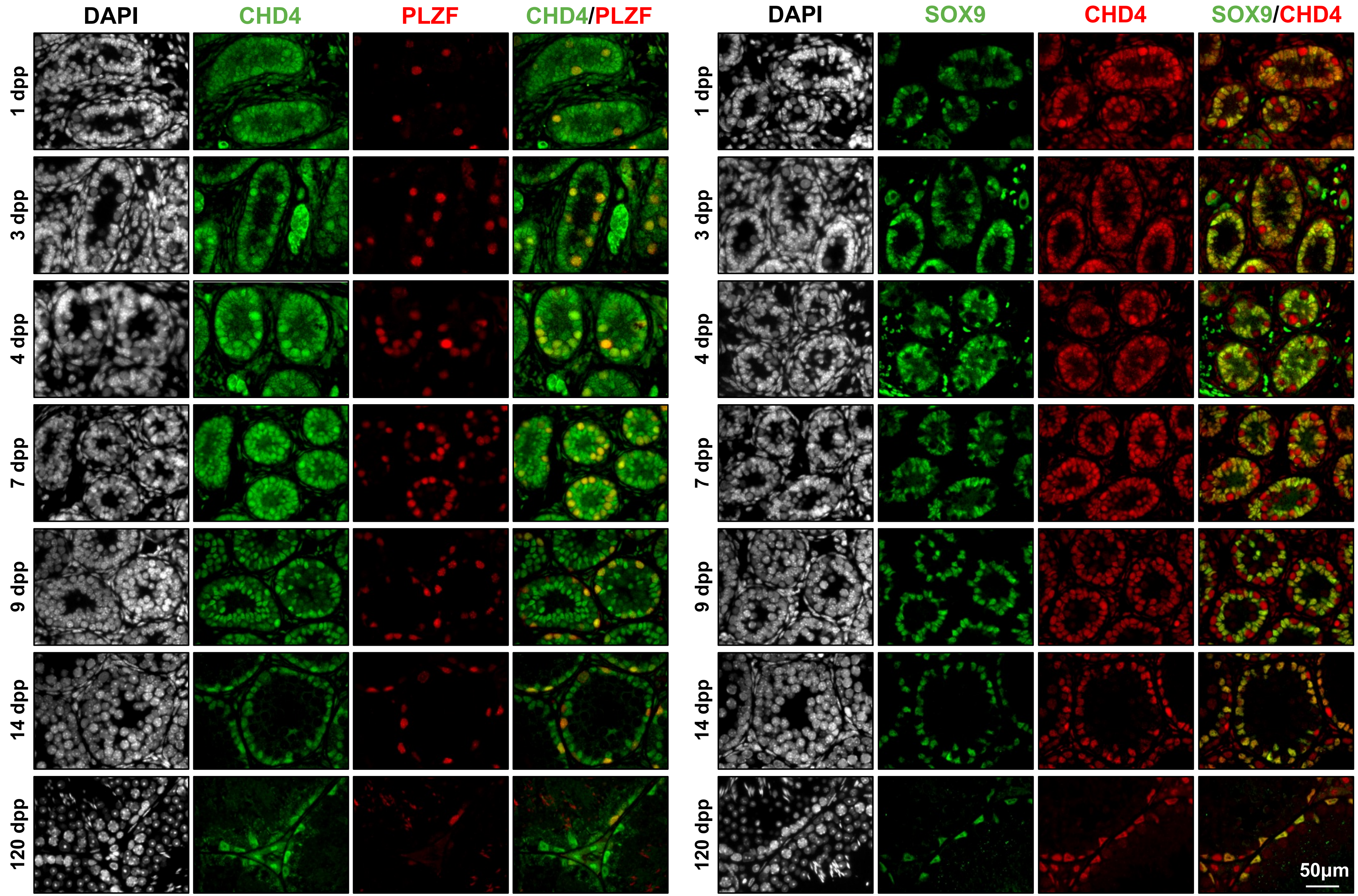

### Supplementary Figure 2

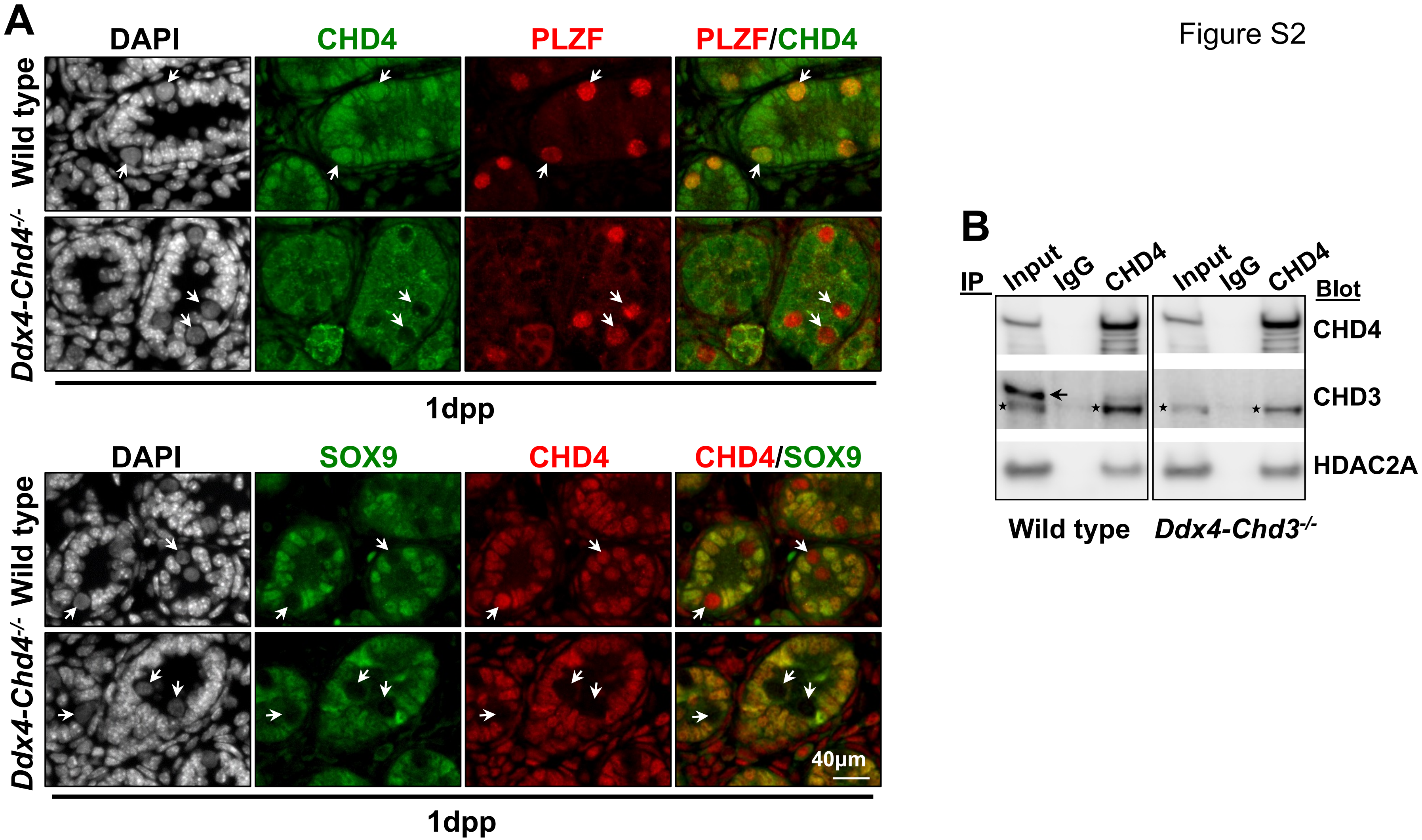

### Supplementary Figure 3

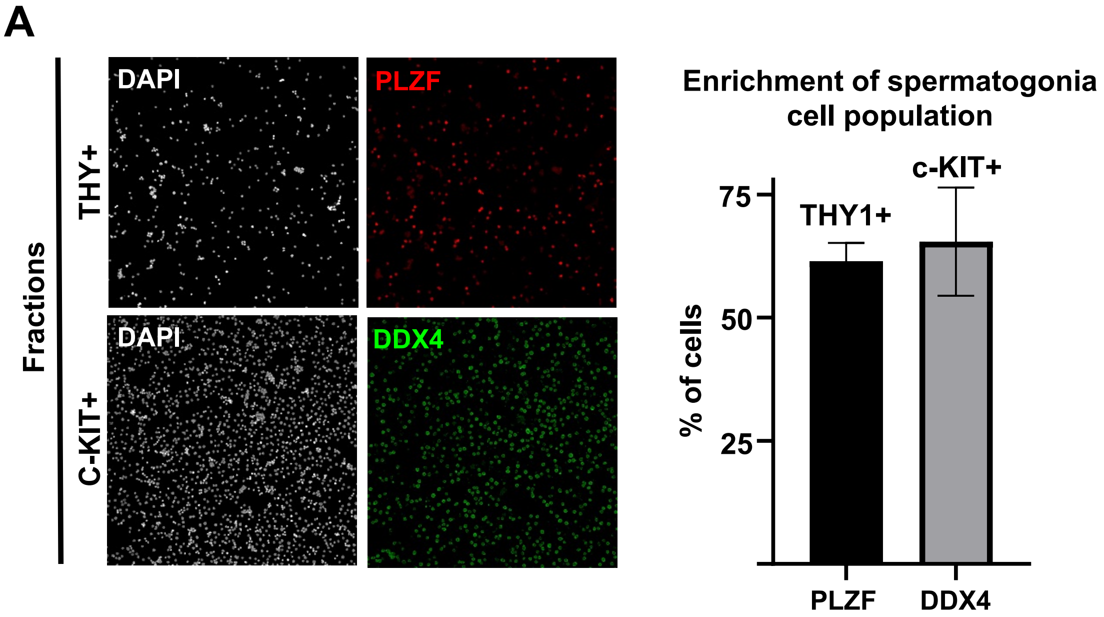

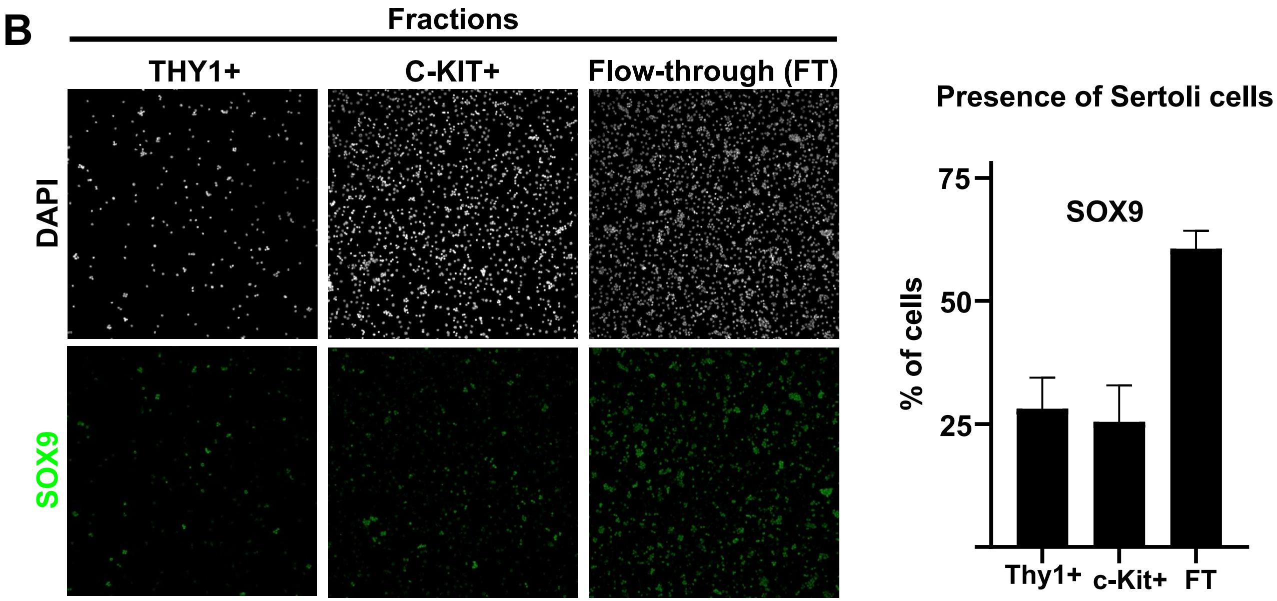

### Supplementary Figure 4

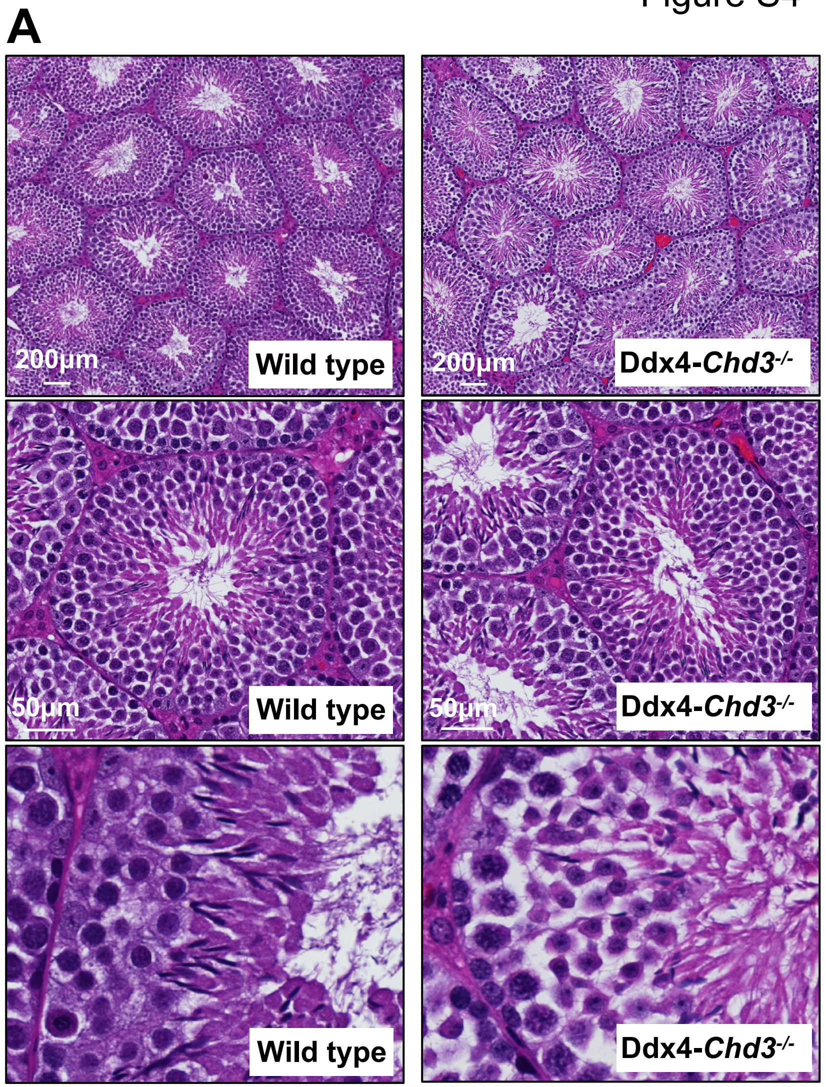

### Supplementary Figure 5

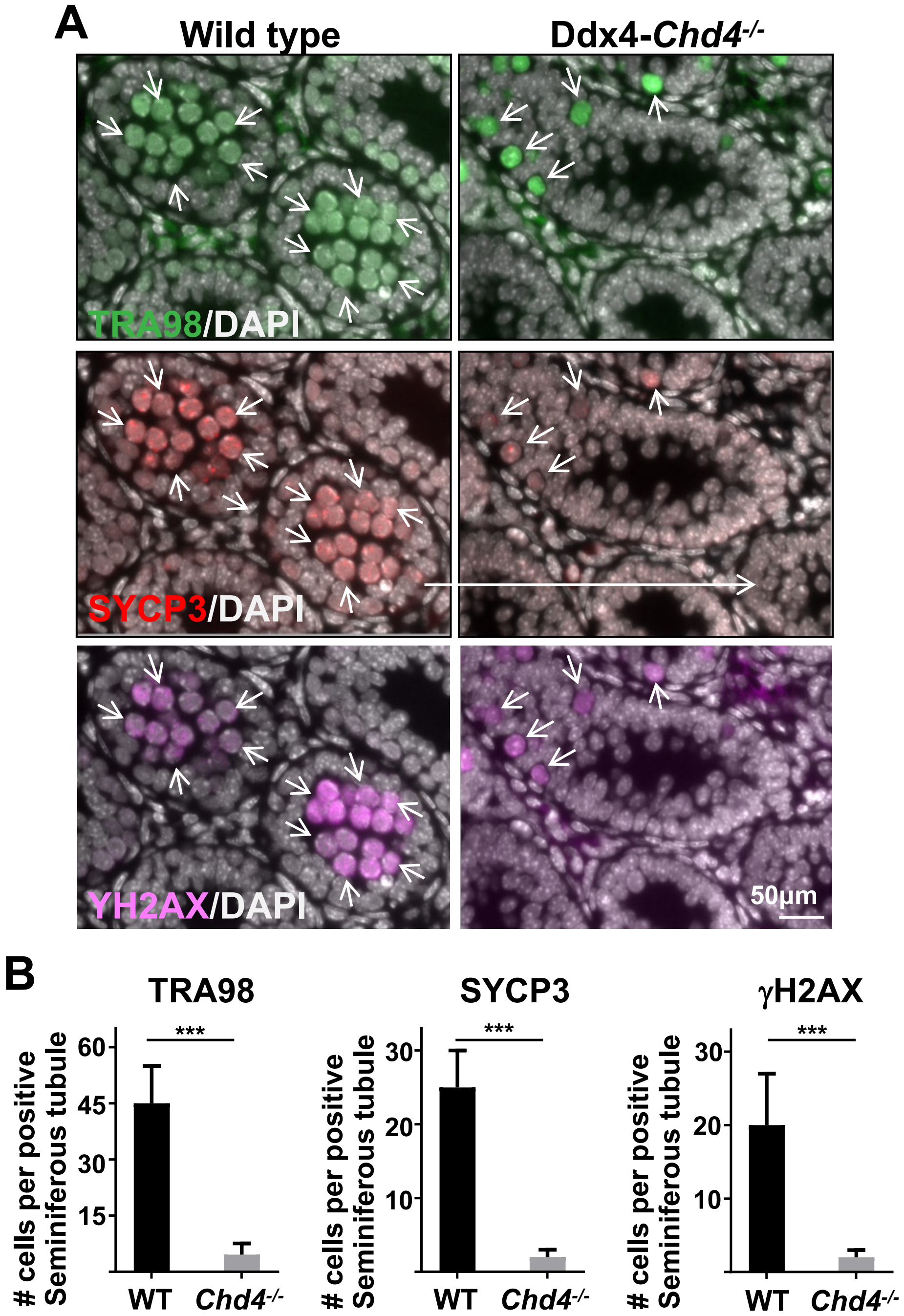

### Supplementary Figure 6

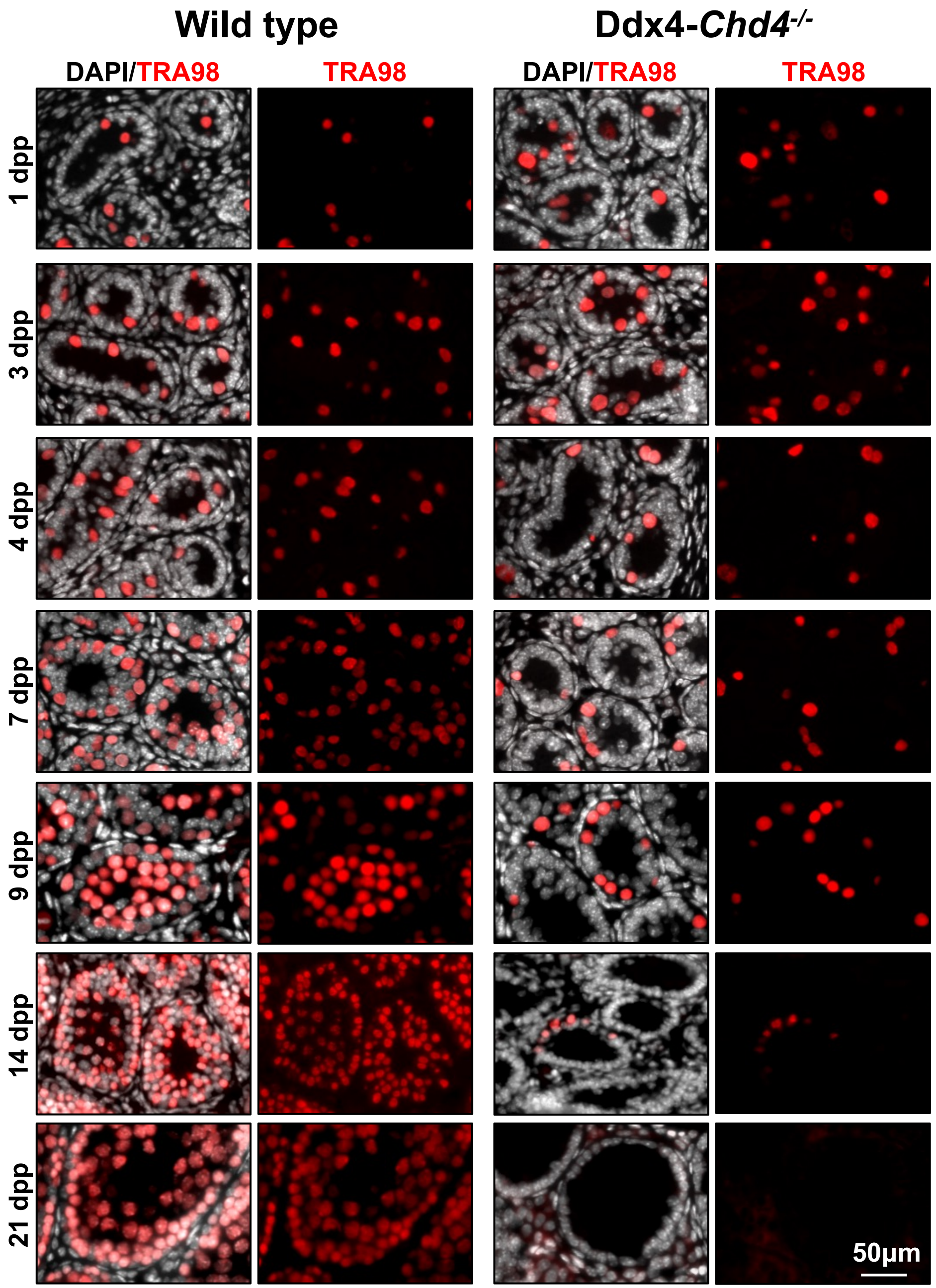

### Supplementary Figure 7

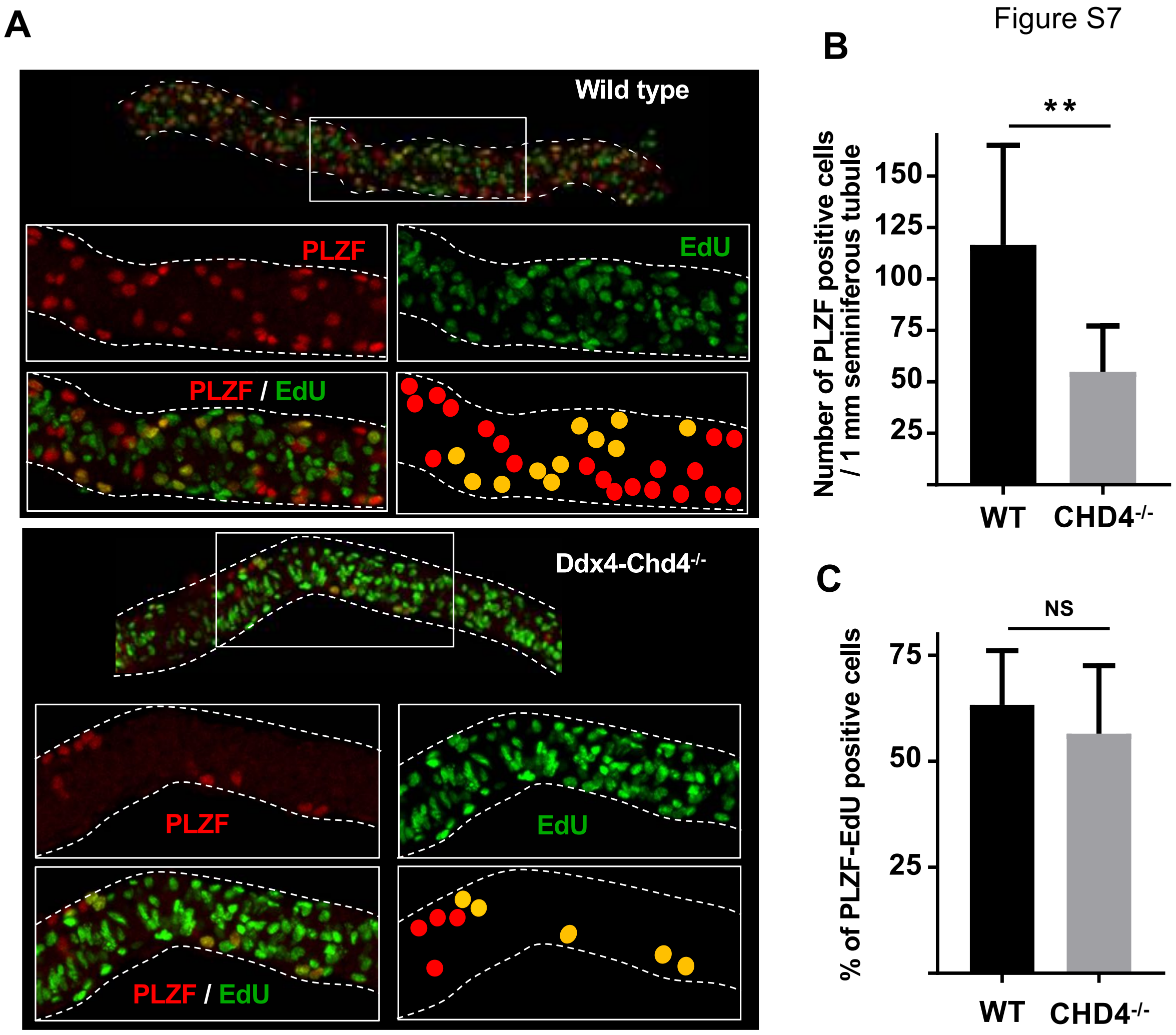
